# ReTrace: Topological evaluation of white matter tractography algorithms using Reeb graphs

**DOI:** 10.1101/2023.07.03.547451

**Authors:** S. Shailja, Jefferson W. Chen, Scott T. Grafton, B.S. Manjunath

## Abstract

We present ReTrace, a novel graph matching-based topological evaluation and validation method for tractography algorithms. ReTrace uses a Reeb graph whose nodes and edges capture the topology of white matter fiber bundles. We evaluate the performance of 96 algorithms from the ISMRM Tractography Challenge and the the standard algorithms implemented in DSI Studio for the population-averaged Human Connectome Project (HCP) dataset. The existing evaluation metrics such as the f-score, bundle overlap, and bundle overreach fail to account for fiber continuity resulting in high scores even for broken fibers, branching artifacts, and mis-tracked fiber crossing. In contrast, we show that ReTrace effectively penalizes the incorrect tracking of fibers within bundles while concurrently pinpointing positions with significant deviation from the ground truth. Based on our analysis of ISMRM challenge data, we find that no single algorithm consistently outperforms others across all known white matter fiber bundles, highlighting the limitations of the current tractography methods. We also observe that deterministic tractography algorithms perform better in tracking the fundamental properties of fiber bundles, specifically merging and splitting, compared to probabilistic tractography. We compare different algorithmic approaches for a given bundle to highlight the specific characteristics that contribute to successful tracking, thus providing topological insights into the development of advanced tractography algorithms.

## 1 Introduction

Tractography is a technique utilized in neuroimaging to reconstruct white matter fiber pathways from diffusion magnetic resonance images (dMRIs). The reconstructed fiber bundles provide valuable insights into the connectivity between different regions of the brain. They play a crucial role in understanding neuroanatomy and studying various brain disorders. For instance, tractography has been used to reveal brain abnormalities across a range of conditions [1, 4, 9], including multiple sclerosis, cognitive disorders, Parkinson’s disease, brain trauma, tumors, and psychiatric conditions. To ensure accurate interpretation of the obtained tractography results, it is vital to evaluate the performance of tractography methods on these neuroanatomical bundles and select appropriate metrics for assessment. Evaluating the performance of tractography methods is a complex task due to the intricate nature of white matter pathways and the challenges associated with capturing their neuroanatomical topology [5, 8, 16].

Empirical studies have examined the effectiveness of many tractography methods in investigating a range of neurodegenerative diseases like Alzheimer’s [21], as well as in measuring patient outcomes following the utilization of tractography for tumor resection [20]. However, the comparison between tractography algorithms remains mainly qualitative for the most part. These studies do not provide a comprehensive evaluation of the tractography algorithms themselves, as they lack a ground truth reference for the reconstructed bundles. This limitation hinders the ability to quantitatively compare standard tracking algorithms and obtain meaningful feedback about their performance.

To quantify tractography, the FiberCup phantom dataset [3] is commonly used. The International Society for Magnetic Resonance in Medicine (ISMRM) organized a tractography challenge [10] on revised FiberCup dataset to establish a ground truth and provide score using the Tractometer [2]. Tractometer provides global connectivity metrics and facilitates extensive assessment of tracking outputs, fiber bundle detection accuracy, and incomplete fiber quantification. As bundle analysis is crucial in neurological studies, the tractograms were divided into 25 major bundles in the ISMRM dataset. To assess the performance of tractography method on bundles, bundle coverage metrics were proposed [10]. These metrics transform the fibers into voxel images, which results in the loss of fiber point-correspondence. They fail to account for many reconstruction errors, thereby yielding inflated and potentially misleading scores. For example, the topological complexity arising due to the geometrical structure and branching within valid bundles is often neglected in voxel-based metrics. In other words, the existing tractography assessment metrics do not answer “how” the fibers are connected but only analyze the connection percentages. Hence, there is a critical need for methods that can quantify the anatomical validity of fiber branching, given the complex fiber topology [19].

Topology pertains to the organization of white matter fibers into structural networks of brain. It considers the fibers’ origination, termination, and branching, as well as their relationship to different brain regions. Neurological disorders can induce topological changes in white matter fibers, such as physical disruptions in cases like stroke, or spatial distortions as seen in brain tumors. To effectively assess the topology of the fiber reconstruction in bundle tracts, we propose ReTrace. It is a novel graph matching algorithm that is based on the construction of a Reeb graph [17, 18]. This graph matching algorithm enables a comprehensive quantitative analysis of topological connectivity patterns by considering both global and local network features. Further, for each quantitative assessment provided by ReTrace, a graph visualization of the bundle in 3D is also associated. This visualization is crucial in analyzing the efficacy of the tractography. Finally, ReTrace can also be tuned to explore the output of a given tractogram in different resolutions. The implementation code and interactive notebooks for utilizing ReTrace are publicly available on GitHub ^3^.

## 2 Tractography evaluation

Quantitative metrics such as global connectivity-dependent metrics in Tractometer [2] and connectivity matrix have been used to evaluate the fiber reconstruction [15]. They give information about the high-level connection of brain regions. However for the assessment of a given bundle tract, they offer a snapshot of valid bundle coverage with their ground truth counterparts and do not account for local topological features. We describe the commonly used metrics in Fig. 1A. While the existing metrics are valuable in assessing volumetric bundle coverage they fall short in capturing fiber continuity, branching, and crossing as highlighted in Fig. 1B, C, and D respectively in the bundle. Some of these limitations are:

**Fig. 1.**
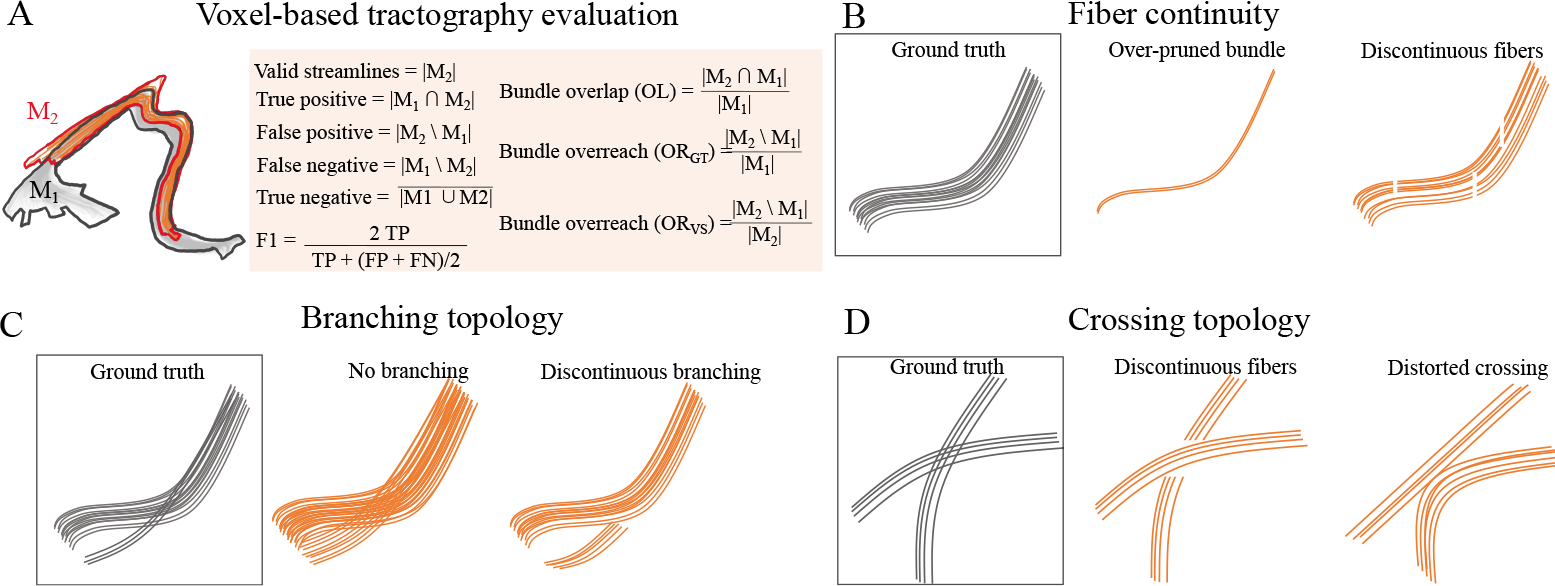
Traditional bundle coverage metrics and their limitations in tractography evaluation. (A) Mask *M*_1_ illustrates the ground truth bundle (black outline), while Mask *M*_2_ reveals valid fibers (red outline) produced by a specific tractography method. Bundle coverage evaluation uses these voxel masks. (B)-(D) display scenarios where the traditional metric falls short. Gray fibers represent the ground truth, while orange fibers depict the reconstructed fibers. (B) highlights potential errors in fiber continuity. (C) demonstrates potential inaccuracies in the branching topology of the tracked fibers, and (D) exposes potential misinterpretations in crossing topology, which despite yielding high scores in bundle coverage, might be anatomically implausible.

1. Voxel-wise agreement between the algorithm’s output and the ground truth is the main emphasis in computing the existing metrics. This overlooks the intricate tractography errors, such as, spurious fibers and incorrect connectivity patterns. Specifically, these metrics often neglect broken fibers as they could still cover the volume despite their discontinuity.
2. Spatial localization is not considered to evaluate the errors between the reconstructed bundles and the ground truth. Similarly, tractography errors cannot be attributed to specific inaccuracies in tracked fibers within the spatial context to guide the design of better algorithms.
3. Branching in the reconstructed pathways is largely inconsequential in the computation of voxel-based metrics. Typically, fiber bundles originate from functional regions, merge into larger pathways for efficiency, and branch out as they approach their respective terminal functional areas. Neuroanatomical studies [5, 17] have pointed out the importance of branching in tractography. But, the conventional metrics (in Fig. 1A) may fail to detect errors related to this branching topology. As such, obscured connections within dense bundles could still achieve high scores despite inaccuracies.
4. Complex fiber orientations such as fiber crossing are not directly captured by the existing tractography evaluation metrics.

Our proposed method addresses these limitations for evaluating tractography methods. It incorporates higher-level tract analysis that considers the topological branching patterns and pathways. Additionally, our method is sensitive to spatial localization, taking into account the precise anatomical locations of inaccuracies within the reconstructed bundles. By addressing the limitations discussed above, our method provides a tunable and robust evaluation of tractography algorithms in terms of accuracy and anatomical fidelity.

## 3 ReTrace: Quantifying topological analysis

ReTrace is an end-to-end tractography evaluation pipeline that starts by processing the given white matter fiber bundles to generate a graph model. Then, it evaluates the quality of fiber reconstruction by comparing two graphs taking into account many topological factors. The pipeline is illustrated in Fig. 2. It is based on the construction of a Reeb graph that provides a topological signature of the fibers. This graph representation makes the tractography evaluation amenable to graph and network theory methods.

**Fig. 2.**
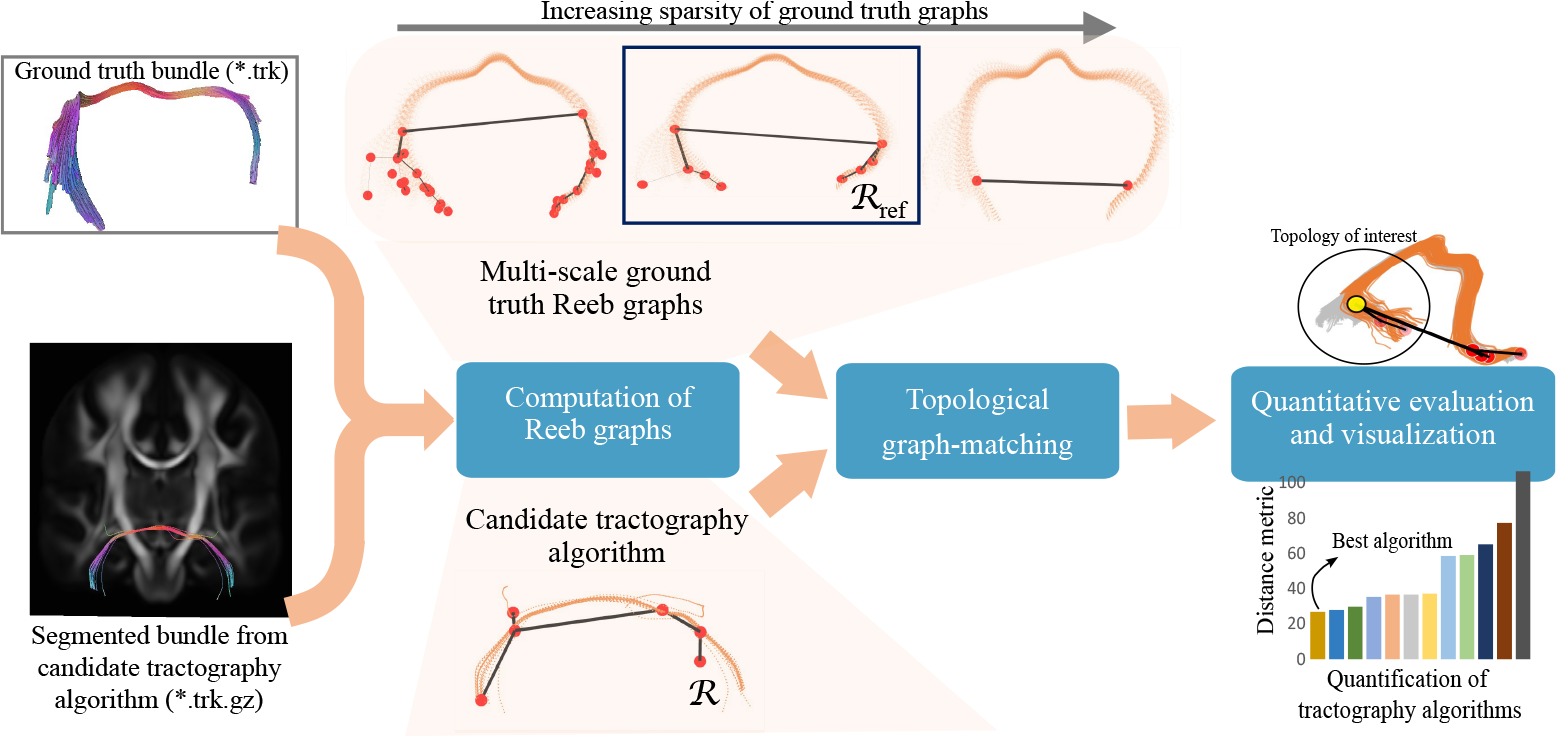
ReTrace evaluates tractography methods both qualitatively and quantitatively by comparing two Reeb graphs, *ℛ* and *ℛ*_ref_. By setting appropriate parameters, Reeb graphs of different resolutions can be computed from the ground truth data. Among the graphs of different sparsity, one is chosen based on the level at which tractography evaluation is desired. Similarly, for the candidate tractography algorithm under evaluation, a Reeb graph is computed from the bundle using the same Reeb graph parameters as the ground truth. The evaluation metric reflects the distance between the two graphs: a higher metric value indicates larger discrepancies, and the candidate algorithm with the lowest metric is assigned the first rank. For qualitative analysis, one can visually examine the Reeb graphs, locating nodes contributing to the distance.

### 3.1 Reeb graphs characterize the topology of anatomical bundles

A Reeb graph representation of white matter fibers has been recently proposed [17, 18]. It provides a concise depiction of trajectory branching structures by constructing undirected weighted graphs. The given bundle tracts (that is, a group of streamlines representing a major bundle) is represented as a graph. The vertices of the graph are critical points where the fibers appear, disappear, merge, or split. Edges of the graph connect these critical points as groups of sub-trajectories of the given bundle. The edge weight is the proportion of fibers that participate in the edge. Three key parameters capture the geometry and topology of fibers:

1. *ε* denotes the distance between a pair of fibers in a bundle, controlling its sparsity. Smaller *ε* values result in denser subtrajectory groups, while larger values allow for sparser groups,
2. *α* represents the spatial length of the bundle, introducing persistence and influencing the extent of bundling, and
3. *δ* determines the bundle thickness to shape the model’s robustness and granularity.

By adjusting these parameters, we can explore the anatomical fiber structure at different scales of sparsity. In this paper, we analyze the reconstruction of anatomical bundles in different spatial resolutions, demonstrating the potential of graph-based tractography validation and evaluation.

*Note:* Throughout the paper, we overlay the Reeb graphs on raw fibers (in orange and ground truth fibers in grey). In the graphs, the nodes are illustrated in red, while edges are shown in black.

### 3.2 Global network features

As discussed, representing the tracts as a graph allows the tractography evaluation to be compatible with existing network and graph theory metrics. For a given Reeb graph *ℛ*(*V, E*), we compute the global network properties that provide a comprehensive view of the network’s structure and behavior. Global features can be as simple as number of nodes (|*V*|), number of edges (|*E*|), or average degree (2*E*/*N*) to more sophisticated metrics such as diameter, assortativity [12], modularity [13], or transitivity depending on the specific application. These network properties provide valuable insights into the structural characteristics of anatomical bundles represented by the Reeb graph. For example, in the Reeb graph, diameter signifies the longest sequence of merging and splitting events along the trajectories. However, global network-based metrics lack specificity and can be challenging to trace back and localize. Therefore, we introduce a novel graph matching algorithm that uses spatial location and provides a quantified distance between the ground truth and computed fiber bundles using local node features.

#### Algorithm 1

Topological Distance Computation

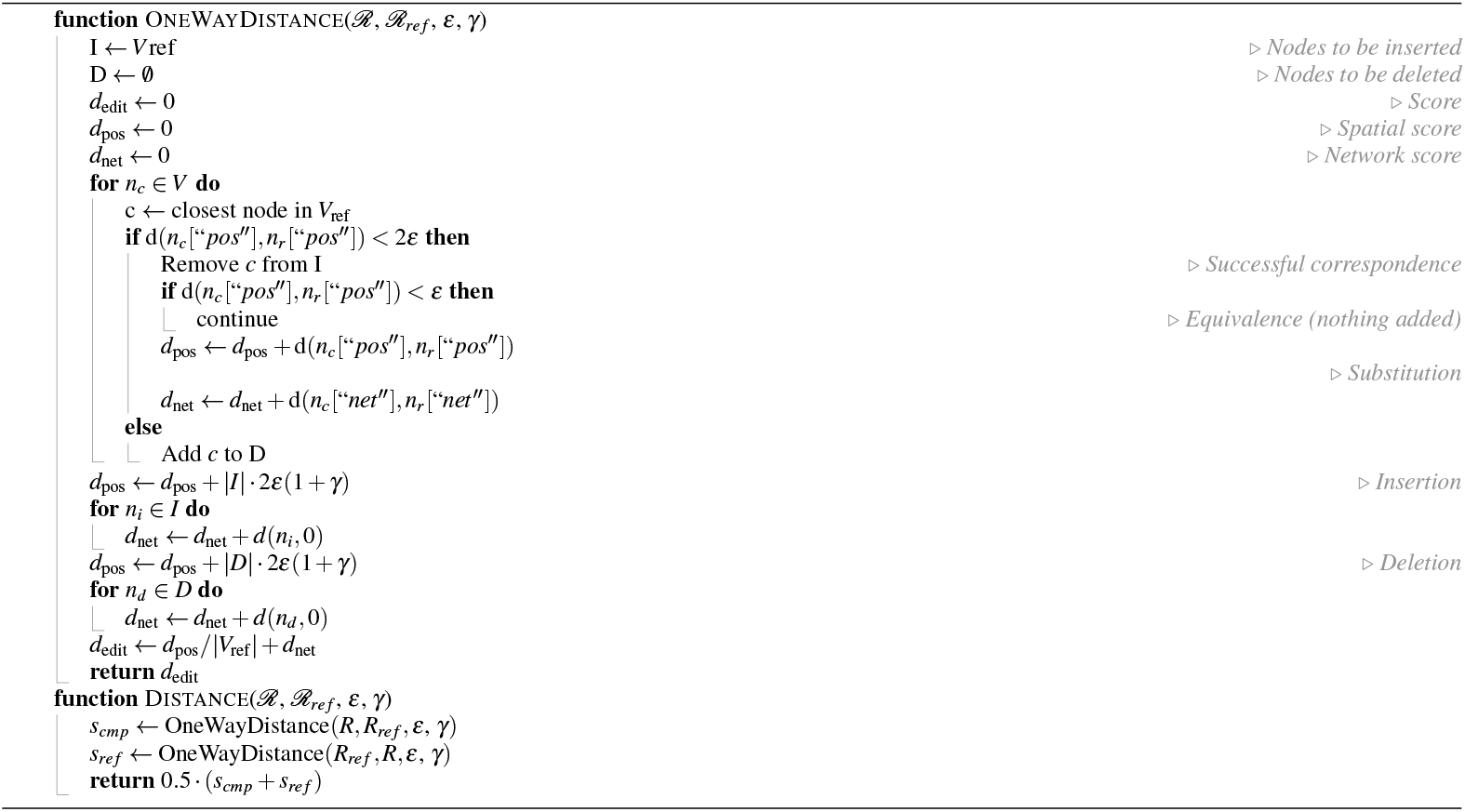

### 3.3 Topological graph matching

For any given node *v* ∈ *V*, we calculate two sets of features — spatial position-based features, denoted as “pos”, and local network-level features, denotes as “net”. Each node of the Reeb graph is linked to its 3D spatial location in the brain, so “pos” corresponds to the 3D coordinate yielding a 3-dimensional feature. On the other hand, the network features are computed using centrality metrics at the node level: degree centrality, closeness centrality, betweenness centrality, and eigenvector centrality [6], producing a 4-dimensional feature.

Our proposed graph matching computation algorithm is adapted from Siminet [11], enhanced to accommodate additional parameters for robustness against noise in tractography and to incorporate network metrics with spatial position as node features. It calculates an edit distance denoted by *d*_edit_ between two Reeb graphs, quantifying the cost of operations needed to transform one graph into another, as outlined in Algorithm 1.

The algorithm begins by iterating through the nodes of the comparison graph *ℛ* and matching each node with its closest spatial counterpart within a search radius. The algorithm determines node correspondences by comparing the spatial distances (“pos”) between nodes and their counterparts against a threshold *ε* (inter-fiber distance used in constructing the Reeb graph). The algorithm also accounts for insertions and deletions by penalizing them with a score *γ* proportional to the Euclidean distance between the centroids of node locations for *ℛ* and *ℛ*_ref_. The proposed metric *d*_edit_ is computed considering the Euclidean distance in node attributes: spatial positions and centrality metrics to compare graph-based representations of white matter bundles, ensuring that layout similarities correspond to similarities in brain regions.

Note that the metric is not normalized, that is, it does not have the standard 0-1 limits. We opt to keep the distances between graphs unbounded to account for relative variations (in some case, the distance could be very large). The ultimate aim of a new tractography design would be to minimize this distance. A potential limitation, or rather a feature, is that the user needs to select the resolution parameters based on the desired comparison of the bundle tracts. While this might appear a manual step, the preset parameters generally serve well as a great starting point for all the major bundle tracts. Also, the performance of all tractography evaluation metrics, both existing and our proposed metric, depends on the quality of bundle segmentation. Hence, further research on bundle segmentation could enhance metric performance.

## 4 Results

In this section, we apply the ReTrace pipeline to both synthetic and real diffusion MRI datasets to demonstrate its efficacy.

### The ISMRM 2015 Tractography Challenge

The challenge presented participants with a clinical-style dataset: a 2mm isotropic diffusion acquisition with 32 gradient directions and a b-value of 1000 s/mm^2^. The task was to reconstruct fiber pathways using a realistically simulated replication of a whole-brain diffusion-weighted MR image. The challenge resulted in 96 tractogram submissions, available publicly for download^4^. These submissions collectively represent a broad range of tractography pipelines, encompassing varied pre-processing, tractography, and post-processing algorithms. This provides a diverse platform for quantitative and qualitative analysis of tractography methods. We assign each algorithm a unique algorithm number, ranging from 1-96 (see the Supplementary Information for the mapping of these ids to the original submission numbers).

For tractography evaluation, we compute the valid bundles for each submission using the ROI-based bundle segmentation system proposed by the challenge organizers [14]. An expert segmented ground truth bundle segmentation provided by the challenge allowed us to assess the submitted tractography methods based on traditionally computed metrics, establishing a baseline for comparison. The extracted bundles were then processed using the ReTrace pipeline. We can fine-tune the computation of Reeb graphs using the robustness parameters (*ε, α*, and *δ*), adjusting the granularity of the desired analysis. In this study, these are set to *ε* = 2.5, *α* = 5, and *δ* = 5. The results for various bundles using our proposed metric compared with the traditional bundle coverage metrics are illustrated in Fig. 3. When the candidate tractography has less than 5 fibers, Reeb graphs are not constructed (automatically with the Reeb graph parameter *δ* = 5) and the algorithm is rejected without rank assignment. For a more dense analysis *δ* may be set to 0. The comparison of existing bundle coverage metrics against our proposed *d*_edit_ metric is presented in Table 1.

**Table 1.** Comparison of *d*_edit_ with existing bundle coverage metrics. The columns labeled as “#” represent the rank 1 algorithm number, while “val” denotes the respective metric values for each metric. Top-ranking methods according to *d*_edit_ differ from those determined by traditional metrics for most bundles. We demonstrate that the reconstruction ranked the best using *d*_edit_ effectively captures branching topology that is overlooked in evaluations using existing methods.

| Bundle | FN | | FP | | OL | | ORGT | | ORVS | | TP | | VS | | Endpoints OL | | Endpoints OR | | fl | | $d_{edit}$ | |
| --- | --- | --- | --- | --- | --- | --- | --- | --- | --- | --- | --- | --- | --- | --- | --- | --- | --- | --- | --- | --- | --- | --- |
|  | # | val | # | val | # | val | # | val | # | val | # | val | # | val | # | val | # | val | # | val | # | val |
| CP | 59 | 1134 | 75 | 38 | 59 | 0.29 | 75 | 0.02 | 32 | 0.42 | 59 | 474 | 56 | 34 | 59 | 14 | 42 | 1 | 59 | 0.33 | 58 | 40.1 |
| CA | 37 | 603 | 60 | 72 | 37 | 0.59 | 60 | 0.05 | 49 | 0.43 | 37 | 876 | 38 | 75 | 43 | 22 | 49 | 2 | 37 | 0.52 | 37 | 7.3 |
| SCP right | 26 | 471 | 44 | 6 | 26 | 0.88 | 44 | 0 | 44 | 0.08 | 26 | 3467 | 57 | 1768 | 57 | 242 | 44 | 1 | 36 | 0.57 | 36 | 10.6 |
| SCP left | 26 | 440 | 34 | 12 | 26 | 0.89 | 34 | 0 | 77 | 0.05 | 26 | 3638 | 57 | 2245 | 25 | 352 | 34 | 0 | 16 | 0.63 | 3 | 10.9 |
| Fornix | 26 | 2698 | 77 | 40 | 26 | 0.75 | 77 | 0 | 77 | 0.07 | 26 | 7971 | 25 | 2595 | 25 | 698 | 77 | 3 | 46 | 0.57 | 18 | 8.7 |

**Fig. 3.**
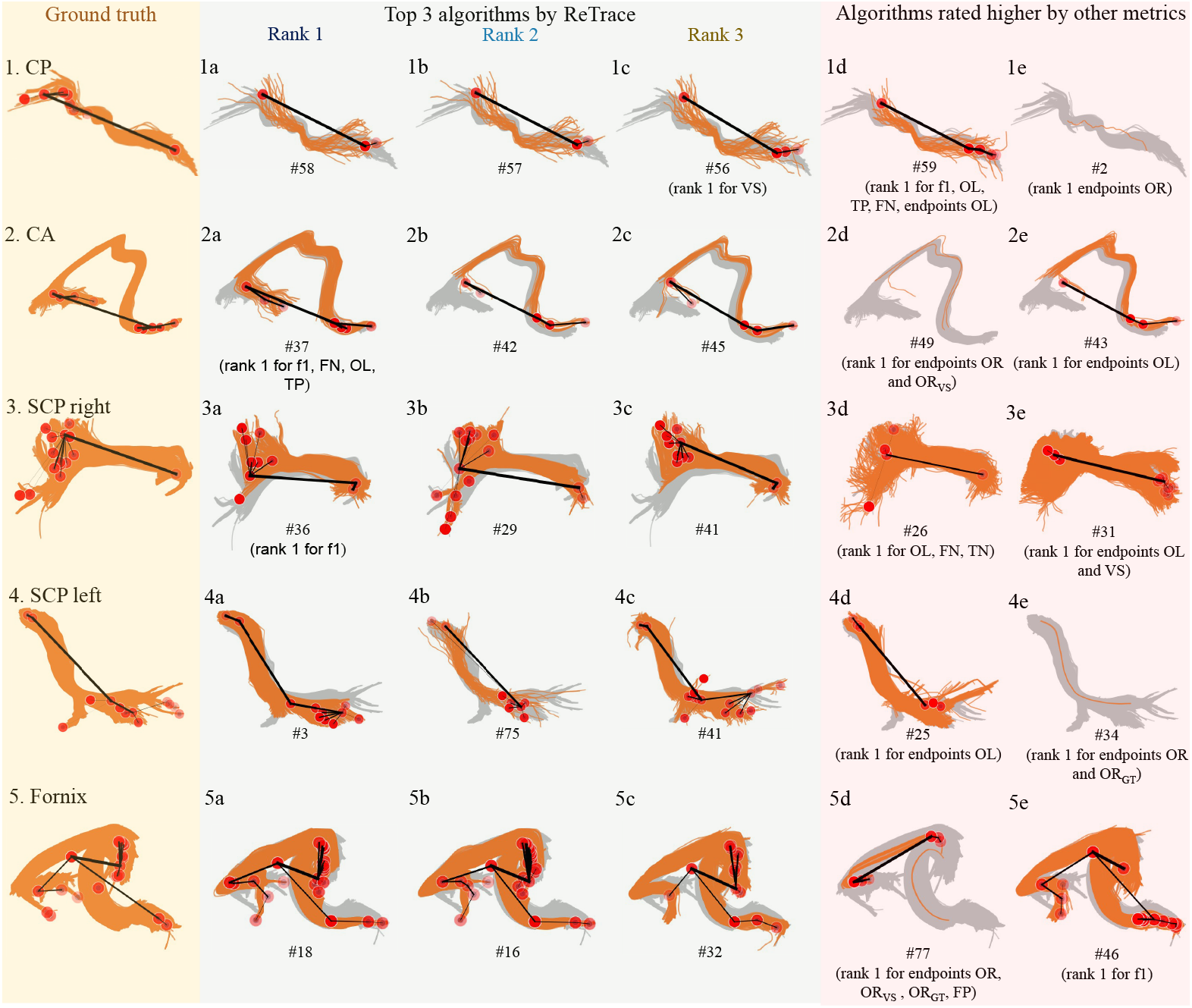
Evaluation of tractography algorithms using ReTrace for bundles in the ISMRM data. The leftmost column displays the ISMRM ground truth fiber bundles with associated Reeb graphs overlaid on the bundles. For the other columns, grey streamlines denote the ground truth, overlaid by orange streamlines representing the candidate tractography algorithm’s results. On top of these, the Reeb graphs are overlaid with red nodes and black edges. The bundles illustrated in the figure are labeled from 1 to 5: CP, CA, SCP_right_, SCP_left_, and Fornix respectively. For each bundle (in a row), the next three columns (a-c) highlight the algorithms that achieved the highest rank according to our proposed metric. Notice that the algorithms that are ranked high on existing metrics (displayed in columns d-e), do not show the best tracked bundle. The reconstructions that do not capture the branches near the end of the bundles are ranked highly by other metrics. Similarly, fiber continuity and distorted fibers in the reconstructed output are given high ranks with existing metrics.

All submissions faced difficulties reconstructing the smaller bundles, such as the anterior (CA) and posterior commissures (CP), which possess a cross-sectional diameter of no more than 2 mm. Due to the minimal branching owing to fewer recovered fibers, ReTrace performs similar to the existing methods on these bundles. However, as the number of fibers increases for bundles like SCP_right_, SCP_left_, and Fornix (as shown in Fig. 3), the importance of accurately branching towards the end of the fibers became apparent. This leads to different algorithms being top-rank with our method as compared with existing metrics. As an example, while algorithm 36 and 29 achieve high ranks for the ReTrace evaluation in SCP_right_, they do not perform as well according to the existing metrics. It is notable that simply increasing fiber count to cover the bundle volume without appropriate branching does not lead to a high ReTrace ranking. This is evident in algorithms 26 and 31 (Fig. 3d and 3e) as they are ranked high according to OL, FN, TN, endpoints OL, and VS metrics, but do not contribute significantly to the ReTrace ranking. The evaluation of all other bundles in the ISMRM dataset using *d*_edit_ along with their interactive visualizations are available on our GitHub repository. Beyond the topological distance that we calculate here, various other graph attributes such as variations in the number of nodes, edges, average degree, and diameter can also be computed and are available in our repository ^5^. They collectively capture different facets of the graphs, and as a consequence, highlight various properties of the tractography reconstruction.

Different tractography methods were employed by different teams. The most important preprocessing, postprocessing, and tractography steps among top-performing algorithms are shown in Fig. 4. For example, CP proved difficult to reconstruct for all algorithms, necessitating the use of extensive denoising and correction methods, as well as probabilistic methods for fiber tracking. Conversely, for other bundles, minimal preprocessing was required and deterministic fiber tracking was sufficient for accurate branching. The insights on the steps of the algorithms offer a topological perspective on designing advanced tractography algorithms and pipelines for future studies. We observe that none of the algorithms perform consistently well for all the bundles. So, an additional advantage of this exploration is that new tractography algorithms may be designed, tailored to specific bundles or neuroanatomical properties of interest.

**Fig. 4.**
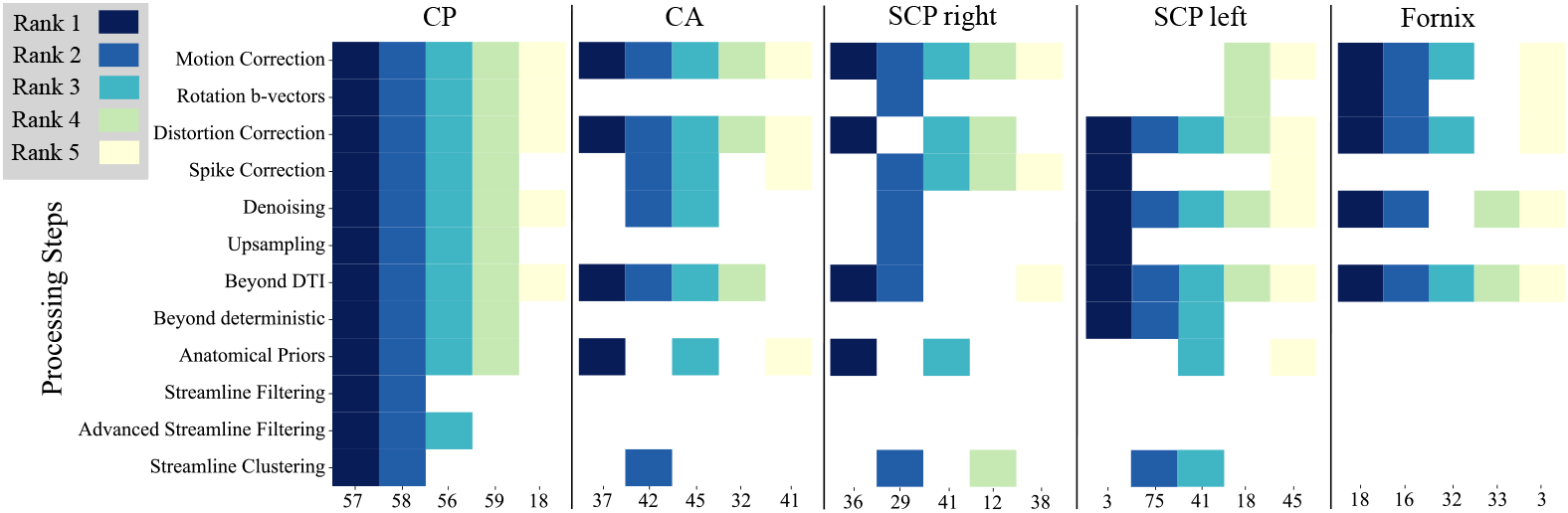
Correlation between processing steps and the successful reconstruction of the major bundles as assessed by Re-Trace. Different colors represent the ranking of methods according to the ReTrace pipeline for five different bundles (rank 1 (dark blue) to rank 5 (yellow)). Y-axis indicate various steps of tractography: Preprocessing (from motion correction to upsampling); Tractography (diffusion modeling beyond DTI such as constrained spherical deconvolution, and tractography methods beyond deterministic approaches such as probabilistic tractography); Postprocessing (incorporation of anatomical priors to streamline clustering). The importance of each step is evident with how prevalent it is in the top ranked algorithms for each bundle.

The results in this paper are in contrast with the ranking of tractography algorithms discussed in the existing literature [10, 14] but also confirm the findings of the importance of various steps in tractography [15]. The runtime of the pipeline depends on the number of fibers and the desired resolution of the Reeb graph. After computing the graphs for a bundle, our evaluation method takes less than 120 seconds to obtain the results on an Intel Xeon CPU E5-2696 v4 @ 2.20 GHz.

### 4.2 Evaluating tractography algorithms in fiber-crossing regions

Tractography’s accuracy is influenced by the number of distinct fiber orientations per voxel, as fiber-crossing regions pose considerable challenges. A probabilistic white matter atlas highlights these areas with high fiber-crossing frequencies [19]. Conventional metrics may not fully capture an algorithm’s ability to accurately resolve these complex regions, instead reflecting its capacity to generate copious or elongated fibers, regardless of their neurological plausibility or accuracy.

For example, a tractography algorithm might generate spurious fibers connecting distinct bundles in crossing fiber scenarios, falsely inflating the fiber count. This inflation could distort performance evaluations if they rely solely on fiber numbers. Endpoint-matching metrics, such as overlap or intersection between reconstructed and ground truth fibers, could also be misleading in the presence of crossing fibers: overlaps may not indicate accuracy if the algorithm incorrectly generates disrupted or distorted fibers due to crossing fibers. Spatial accuracy metrics, like the average or Hausdorff distance [7] between reconstructed and ground truth fibers, could also be skewed by crossing fibers. These regions may compromise the algorithm’s capability to accurately resolve individual bundles, resulting in spatial inaccuracies or increased distances.

#### HCP Dataset

The efficacy of ReTrace is not limited to synthetic data alone. To demonstrate its applicability to real datasets and to show how Retrace handles fiber crossing, we use the average HCP 1065 template, constructed from the diffusion MRI data of 1065 subjects from the Human Connectome Project (HCP)^6^. We use the 1 mm population-averaged FIB file in the ICBM152 space for fiber tracking in DSI Studio.

ReTrace handles fiber crossings effectively, as demonstrated in Fig. 5. We select a small region of interest (a 2mm isotropic 3D region) from the probabilistic atlas in an area with a high probability (∼ 0.9) of double-crossing fibers. We use the deterministic streamline tracking method implemented in DSI Studio^7^ to compute the fibers with the parameters set (angular threshold, step size, min length, max length, terminate if seeds, iterations for topological pruning) to 35, 1, 70, 200, 1000, and 16, respectively. This allowed us to observe successful tracking without broken or distorted fibers. To mimic the spurious broken fibers that tractography methods may yield, we set the parameters to 35, 1, 0, 100, 1000, and 16. For observing the angular distortion where the fiber bends and follows a different path, we set the parameters as 90, 1, 70, 100, 1000, and 16. The resulting Reeb graphs clearly highlight how their nodes capture successful and unsuccessful tracking. Nodes formed near intersections indicate broken or bent fibers, as shown in Fig. 5. Whenever fibers travel together in a group, they form an edge in the graph. Any alteration within this group prompts a critical event, resulting in a node in the Reeb graph. Consequently, if a fiber breaks or diverges, the associated group changes, generating a node. This node, present at the merging point, contributes to a larger distance value. In the topological distance computation, *d*_edit_, this node could be weighted more if the goal is to assess an algorithm’s tracking ability in ambiguous fiber orientations. By providing a 3D location for our algorithm’s attention, any discrepancy within that location can significantly affect the overall *d*_edit_ computation. The code is open source and can be tailored to specific needs.

**Fig. 5.**
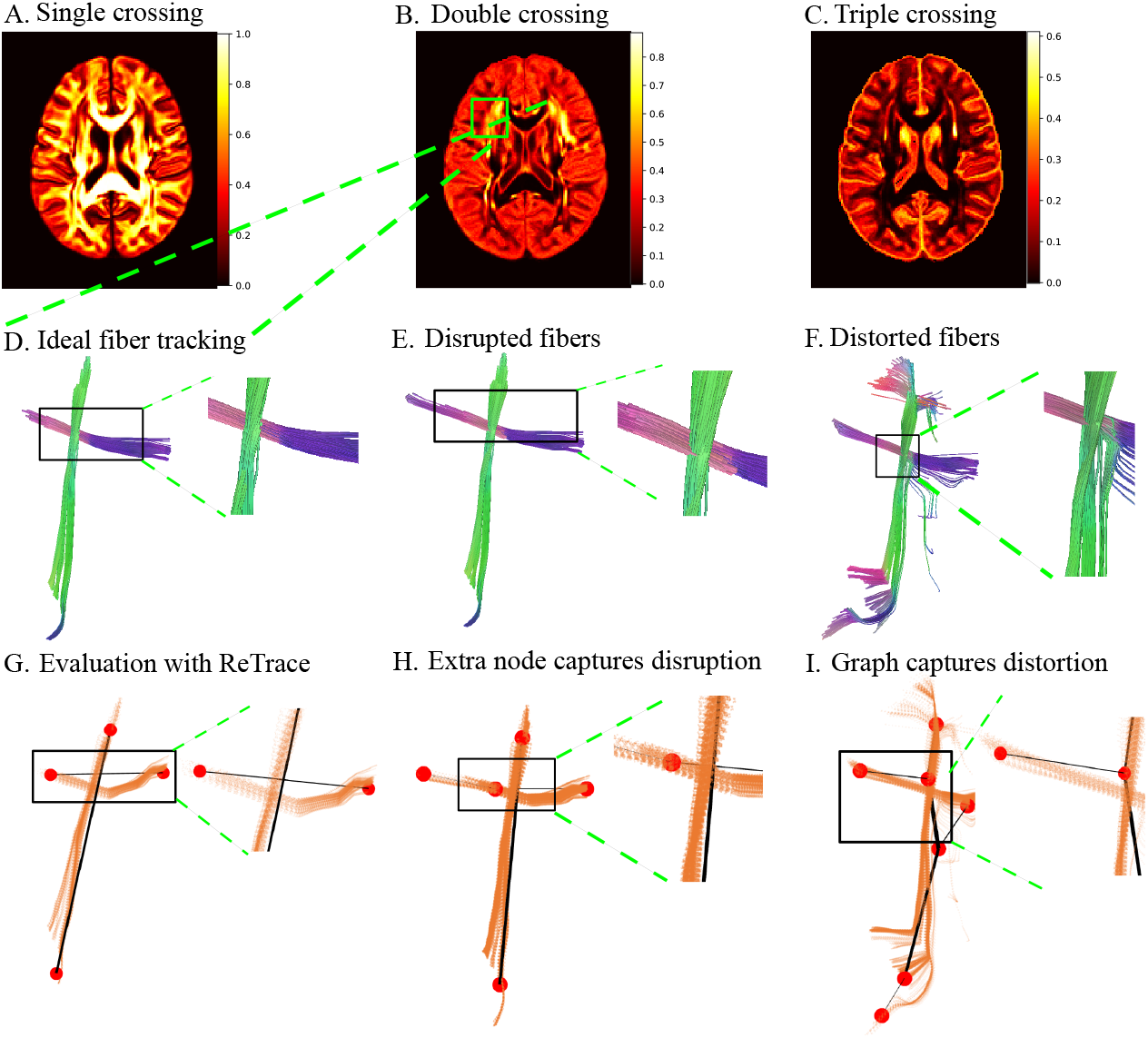
Evaluation of a tractography algorithm for fiber crossing. (A)-(C) show the center slice of the probabilistic directions atlas. This atlas indicates single, double, or triple crossing by the probability of the presence of diffusion tensor oriented in one, two, or three directions. A small ROI with high double crossing fiber probability (shown in green) is selected. (D)-(F) demonstrate the reconstruction of fibers using the streamline method implemented in DSI Studio. (D) shows successful tracking of fiber crossings with ideal parameters. (E) shows fibers that terminate abruptly near the crossing as a result of reconstruction with different parameters. (F) shows bending or distortion of fibers when encountering multiple diffusion orientations. (G)-(I) present the constructed Reeb graphs overlaid on the computed fibers for each case, respectively (shown below each reconstruction). An additional node is observed in the Reeb graph that captures the fiber crossing or bending. A thicker edge in (F) indicates bending of the fibers, whereas ideally, the fiber should only cross without bending.

## 5 Conclusion

This paper introduces ReTrace, an innovative evaluation method for tractography algorithms. This method addresses existing metrics’ limitations by focusing on the topological accuracy of reconstructed pathways. We applied ReTrace to both synthetic and real-world datasets to demonstrate the branching fidelity and errors such as broken or bent fibers in tractography reconstruction. With the ISMRM dataset, we ranked 96 algorithms on different neuroanatomical bundles. The rankings proposed by our method are in contrast with the rankings using the conventional voxel-based tractography metrics. We discuss this difference in ranking and performance of different algorithms by highlighting the topological features such as branching, fiber continuity, localization, and crossing. We demonstrated the utility of our method on the HCP dataset where we show the importance of reconstructed fiber crossing and discuss the performance of standard tractography algorithms. Our approach is disease-agnostic, does not require brain registration to an atlas, and works across different acquisition protocols. It is important to note that bundle segmentation is a common bottleneck in any evaluation metric. Therefore, advancements in segmentation research could greatly enhance these evaluations. The results on tractography comparison presented here could be extended to be used as a cost function for data-driven machine learning methods, like generative adversarial networks. With feedback from neuroscientists, we hope that the results in this paper will pave the way forward in improving existing tractography methods.

## 6 Acknowledgement

The authors acknowledge Vikram Bhagavatula for his assistance in a preliminary implementation of the edit distance. Data were provided by The ISMRM 2015 Tractography Challenge and by the Human Connectome Project, WU-Minn Consortium (Principal Investigators: David Van Essen and Kamil Ugurbil; 1U54MH091657) funded by the 16 NIH Institutes and Centers that support the NIH Blueprint for Neuroscience Research; and by the McDonnell Center for Systems Neuroscience at Washington University.

## Footnotes

3 https://github.com/s-shailja/ReTrace

4 https://zenodo.org/record/840086

5 https://github.com/s-shailja/ReTrace

6 https://brain.labsolver.org/hcp_template.html

7 https://dsi-studio.labsolver.org/

