## Supplementary Information for "ReTrace: Topological evaluation of white matter tractography algorithms using Reeb graphs"

Mapping of algorithm IDs to the original submission IDs:

| Submission ID | Algorithm ID |
|---|---|
| 1_0 | 1 |
| 1_1 | 2 |
| 1_2 | 3 |
| 1_3 | 4 |
| 1_4 | 5 |
| 2_0 | 6 |
| 3_0 | 7 |
| 3_1 | 8 |
| 3_2 | 9 |
| 3_3 | 10 |
| 3_4 | 11 |
| 4_0 | 12 |
| 5_0 | 13 |
| 5_1 | 14 |
| 6_0 | 15 |
| 6_1 | 16 |
| 6_2 | 17 |
| 6_3 | 18 |
| 6_4 | 19 |
| 7_0 | 20 |
| 7_1 | 21 |
| 7_2 | 22 |
| 7_3 | 23 |
| 8_0 | 24 |
| 9_0 | 25 |
| 9_1 | 26 |
| 9_2 | 27 |
| 9_3 | 28 |
| 9_4 | 29 |
| 10_0 | 30 |
| 10_1 | 31 |
| 10_2 | 32 |
| 10_3 | 33 |
| 10_4 | 34 |
| 10_5 | 35 |
| 10_6 | 36 |
| 10_7 | 37 |
| 10_8 | 38 |
| 10_9 | 39 |
| 10_10 | 40 |
| 10_11 | 41 |
| 10_12 | 42 |
| 10_13 | 43 |
| 10_14 | 44 |
| 10_15 | 45 |
| 10_16 | 46 |
| 10_17 | 47 |
| 10_18 | 48 |
| 10_19 | 49 |
| 11_0 | 50 |
| 11_1 | 51 |
| 12_0 | 52 |
| 12_1 | 53 |
| 12_2 | 54 |
| 12_3 | 55 |
| 13_0 | 56 |
| 13_1 | 57 |
| 13_2 | 58 |
| 13_3 | 59 |
| 14_0 | 60 |
| 14_1 | 61 |
| 14_2 | 62 |
| 15_0 | 63 |
| 16_0 | 64 |
| 16_1 | 65 |
| 16_2 | 66 |
| 16_3 | 67 |
| 16_4 | 68 |
| 17_0 | 69 |
| 17_1 | 70 |
| 17_2 | 71 |
| 17_3 | 72 |
| 17_4 | 73 |
| 18_0 | 74 |
| 18_1 | 75 |
| 18_2 | 76 |
| 18_3 | 77 |
| 18_4 | 78 |
| 19_0 | 79 |
| 19_1 | 80 |
| 19_2 | 81 |
| 20_0 | 82 |
| 20_1 | 83 |
| 20_2 | 84 |
| 20_3 | 85 |
| 20_4 | 86 |
| 20_5 | 87 |
| 20_6 | 88 |
| 20_7 | 89 |
| 20_8 | 90 |
| 20_9 | 91 |
| 20_10 | 92 |
| 20_11 | 93 |
| 20_12 | 94 |
| 20_13 | 95 |
| 20_14 | 96 |
